# Systematic Functional Annotation and Visualization of Biological Networks

**DOI:** 10.1101/030551

**Authors:** Anastasia Baryshnikova

## Abstract

Large-scale biological networks map functional connections between most genes in the genome and can potentially uncover high level organizing principles governing cellular functions. These networks, however, are famously complex and often regarded as disordered masses of tangled interactions (“hairballs”) that are nearly impenetrable to biologists. As a result, our current understanding of network functional organization is very limited. To address this problem, I developed a systematic quantitative approach for annotating biological networks and examining their functional structure. This method, named Spatial Analysis of Functional Enrichment (SAFE), detects network regions that are statistically overrepresented for a functional group or a quantitative phenotype of interest, and provides an intuitive visual representation of their relative positioning within the network. By successfully annotating the *Saccharomyces cerevisiae* genetic interaction network with Gene Ontology terms, SAFE proved to be sensitive to functional signals and robust to noise. In addition, SAFE annotated the network with chemical genomic data and uncovered a new potential mechanism of resistance to the anti-cancer drug bortezomib. Finally, SAFE showed that protein-protein interactions, despite their apparent complexity, also have a high level functional structure. These results demonstrate that SAFE is a powerful new tool for examining biological networks and advancing our understanding of the functional organization of the cell.

## INTRODUCTION

Understanding the functional organization of living cells is essential for predicting their behavior in normal and diseased conditions, and designing effective therapeutic strategies to control it. Despite rapid advancements in technology development and data collection, our current knowledge about functional organization is limited to individual molecular pathways and a few well-characterized processes, whereas the cell as a whole remains a largely obscure system.

A unique opportunity to elucidate the organization of a cellular system comes from budding yeast *Saccharomyces cerevisiae* which benefits from the availability of extensive molecular interaction networks that map physical, biochemical and phenotypic relationships between nearly all genes in the genome (Gavin et al., 2006; Krogan et al., 2006; Zhu et al., 2007; Tarassov et al., 2008; Yu et al., 2008; Mo et al., 2009; Costanzo et al., 2010). Since each type of interaction captures a different aspect of a gene-gene relationship, it is ultimately the integration of all networks that will provide the most comprehensive view of the functional organization of the yeast cell. However, to devise good strategies for integrating networks, we must first understand their individual functional structures, i.e. we must determine which biological functions are represented in each network, which parts of the network these functions are associated with and how they are related to one another. This process, analogous to understanding the functional structure of a newly sequenced genome, can be referred to as functional annotation of biological networks.

Early attempts to visualize and annotate networks, carried out through mostly manual analysis, unveiled their potential for advancing our knowledge about the cell (Costanzo et al., 2010). For example, close examination of the yeast genetic interaction network revealed that genes naturally assembled into 13 major functional groups whose relative positioning within the network was largely consistent with their functional relatedness (Costanzo et al., 2010). To confirm these trends quantitatively and explore the functional structure of other molecular networks, we need to approach network annotation systematically, using rigorous statistical methods and reproducible workflows.

In its simplest form, systematic annotation of a biological network requires three steps. First, we must obtain a comprehensive map of the network showing all of its nodes and their connections to one another. This map can be produced using a network layout algorithm that embeds the network in two- or three-dimensional space and positions all nodes based on their connectivity (Kobourov, 2012). Second, we need to secure multiple independent sources of functional information that would characterize all nodes relative to one another based on a variety of different parameters (e.g., cellular localization and abundance, transcriptional response to a perturbation, mutant phenotype). Such functional resources are readily available in yeast, thanks to the development of numerous genomic assays and a long history of literature curation (Botstein and Fink, 2011). Finally, we must implement automated statistical procedures to overlay functional data onto the network map and locate functionally coherent regions. While functional regions have been discovered on a case-by-case basis (Khatri et al., 2012; Mitra et al., 2013), no method currently exists to identify them exhaustively. Moreover, no method is currently equipped to determine the positioning of functional regions relative to each other and create a functional map of the network that is comprehensive, quantitative and intuitive to biologists, all of which are essential requirements for its correct interpretation.

In theory, some of the existing approaches could be stretched to produce such a functional map. However, doing so would require extensive modifications and/or additional post-processing steps that would drastically reduce the scalability of these approaches for automation. For example, several network algorithms have been developed to search for sets of interconnected nodes that share a common phenotype or a consistent response across experimental conditions (Ideker et al., 2002; Mitra et al., 2013). The main purpose of these algorithms is to evaluate experimental datasets and identify the most promising candidate genes in which phenotype similarity is supported by network connectivity. Since networks themselves are not the focus of the analysis but only independent supporting evidence for a hypothesis, these methods must undergo significant changes in order to be applicable for comprehensive annotation tasks. Similarly, network clustering algorithms could potentially be used to identify sets of densely connected nodes that correspond to known, as well as novel, functional modules (Newman, 2006). However, this strategy disregards loosely connected nodes, causing many sparse yet functionally coherent network regions to remain unannotated. In addition, clustering algorithms partition the network into discrete and, in some cases, overlapping subnetworks, which must be annotated separately and integrated back together to provide a global functional view of the network. Since rapid and reproducible integration of functional annotations has yet to be achieved systematically, the use of clustering algorithms for annotating biological networks is currently impractical. More recently, a spatial statistics-based approach has been proposed to identify functional groups that co-cluster within the network more closely than expected by random chance (Cornish and Markowetz, 2014). While significant co-clustering does indicate that the functional group is strongly associated with the network, it does not reveal in which part of the network the association occurs and what other functional groups are associated nearby. As a result, both coverage and specificity of functional associations cannot be assessed by the co-clustering method and a true functional map of the network cannot be constructed.

To address these and other limitations of the existing methods, I developed an automated procedure for annotating biological networks and generating quantitative and intuitive functional maps. This method, named Spatial Analysis of Functional Enrichment (SAFE), visualizes the network in two-dimensional space and detects network regions that are statistically overrepresented for a functional group or a quantitative phenotype of interest, such as a Gene Ontology (GO) term or a growth response to a chemical treatment. By mapping a function (or a phenotype) to a specific part of the network, SAFE provides statistical evidence, as well as an intuitive visual representation, for the relative positioning of different functions within the network. Using SAFE, I performed an automated functional analysis of the yeast genetic interaction similarity network and showed that an iterative annotation of the network with multiple independent functional resources provides novel insight into the yeast response to chemical treatment. Furthermore, I showed that SAFE can detect functional structure within extremely dense and visually challenging biological networks, such as the extensive yeast protein-protein interaction network. These results indicate that SAFE is a powerful tool for annotating networks in a systematic and unbiased way and a unique framework for investigating the global functional organization of the cell.

## RESULTS

### Description of the SAFE method

SAFE annotates a biological network by calculating and visually representing local enrichment for a set of functional attributes (**Figure 1**; Experimental Procedures). To achieve this, SAFE first generates a two-dimensional map of the network by applying a force-directed network layout algorithm (Experimental Procedures) or adopting a network embedding computed by a third-party software (e.g., Cytoscape (Shannon et al., 2003)) (**Figure 1A**). In a force-directed network layout, connected nodes attract each other, whereas disconnected nodes are pushed apart. As a result, the position of every node is determined by an equilibrium of forces that reflects network topology.

**Figure 1.**
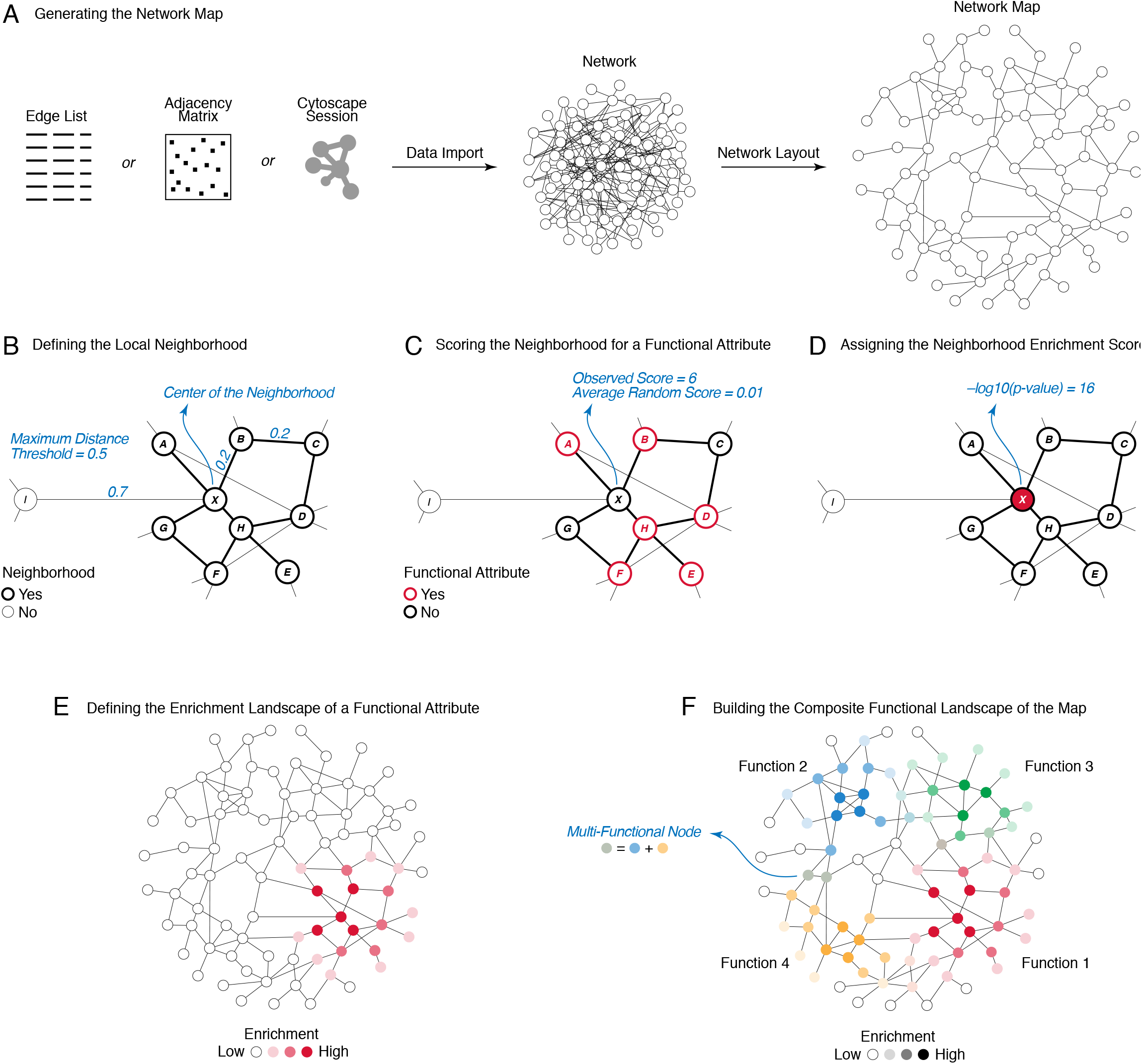
Spatial Analysis of Functional Enrichment (SAFE). For details, see Experimental Procedures. (**A**) Given a biological network, provided as an edge list or an adjacency matrix, SAFE generates a two-dimensional map of the network by applying a force-directed network layout algorithm. Alternatively, SAFE can import a network map directly from the network visualization software Cytoscape (Shannon et al., 2003). (**B**) For each node *X* on the network map, SAFE defines the local neighborhood of *X* by identifying all other nodes (*A*-*H*) located within a chosen distance from it. By default, distance between two nodes is measured by their map-weighted shortest path length (MSPL); however, other distance measures are also available. (**C**) For each neighborhood, SAFE calculates the sum of the neighbors’ values for a functional attribute of interest (e.g., binary annotations to a GO term or quantitative expressions of a phenotype). The neighborhood sum is compared to random expectation and a p-value is calculated. (**D**) The p-value, corrected for multiple testing, is log-transformed into a neighborhood enrichment score and assigned to the center of the neighborhood (node *X*). (**E**) The neighborhood enrichment scores for any given attribute across all nodes in the network define the attribute’s enrichment landscape, which illustrates the distribution of the attribute throughout the network and measures the strength of their association. Color intensity is used to represent variations in enrichment scores. (**F**) Attributes are assigned different colors and summed proportionally to their relative enrichment within each neighborhood. As a result, pure colors indicate regions with a single predominant function, whereas hybrid colors correspond to multi-functional regions. Quantitative enrichment values, color assignments and node-attribute associations are provided as part of SAFE’s output.

For every node on the map, SAFE defines a local neighborhood, which corresponds to the set of nodes located within a chosen distance from it (**Figure 1B**). While several distance metrics are available (Experimental Procedures), the default option is the map-weighted shortest path length (MSPL). According to this metric, distance between two nodes is determined by the length of the shortest path between them, where a path is a set of edges and the length of each edge is its physical extent on the network map (**Figure 1B**). As a result, a node’s neighbors may be directly connected to it or may be a few steps away, provided that the total length of these steps does not exceed the maximum distance threshold (**Figure 1B**).

For every neighborhood, SAFE calculates a set of quantitative scores, each corresponding to the sum of the neighbors’ values for a functional attribute under examination (**Figure 1C**; Experimental Procedures). Attributes may include binary annotations to pre-defined functional groups, such as GO terms, or quantitative phenotypes derived from systematic genomic experiments. To estimate the significance of a neighborhood’s score for a given attribute, SAFE computes a hypergeometric p-value (for binary annotations) or an empirical p-value (for quantitative annotations) based on 1,000 network randomizations that preserve network topology but reshuffle attribute assignments (**Figure 1C**). A log enrichment score (–log_10_ *p*, where *p* is the neighborhood enrichment p-value, corrected for multiple testing) is then assigned to the center of the neighborhood (**Figure 1D**; Experimental Procedures).

The enrichment scores for any given attribute across all nodes on the map define the attribute’s enrichment landscape (**Figure 1E**), which has a characteristic size (i.e., the number of enriched neighborhoods), shape (i.e., the area of the map covered by the enriched neighborhoods) and relief (i.e., the peaks and valleys in the enrichment). Through these properties, the enrichment landscape reflects the distribution of the attribute across the map and measures the strength of its association with the network.

To combine multiple enrichment landscapes into a single functional map of the network, SAFE pairs each attribute with a distinct color and sums the colors proportionally to the relative enrichment of their corresponding attributes within each neighborhood (**Figure 1F**). As a result of this operation, a peak of color intensity indicates that the region is dominated by a single functional attribute, whereas hybrid colors are representative of multi-functional regions (**Figure 1F**). This approach produces a comprehensive view of all functional attributes and a quantitative representation of their enrichments relative to one another.

If several attributes are closely related and map to the same region of the network (as occurs, for example, for a GO term and its descendants), SAFE can combine them into a functional domain based on the similarity of their enrichment landscapes. To facilitate the interpretation of the resulting functional map, all attributes within a domain are assigned the same color and labeled by a single tag list, composed of the five most recurrent words among the attributes’ names (Experimental Procedures).

### SAFE annotates the yeast genetic interaction similarity network with GO terms

To test the SAFE method, I used Gene Ontology biological process terms as attributes to annotate the yeast genetic interaction similarity network (Costanzo et al., 2010) (**Figure 2**). A genetic interaction is a phenotypic relationship between two genes that occurs when the phenotype of the double mutant deviates from the expected combination of the phenotypes of the two single mutants (Baryshnikova et al., 2013). Genetic interactions are strongly associated with genes sharing a common biological function or acting within the same biological process: these genes often genetically interact with each other and show similar genetic interactions with other genes (Baryshnikova et al., 2013). Similarity of genetic interaction profiles is a particularly strong predictor of functional relationship and has enabled the construction of a functional network connecting ∼75% of all genes in the yeast genome (Baryshnikova et al., 2010b; Costanzo et al., 2010). A high-confidence version of this network, consisting of 2,838 nodes and 10,016 edges (**Figure 2A**), was visualized and manually annotated in its original study (Costanzo et al., 2010) and therefore provides a good test case for SAFE.

**Figure 2.**
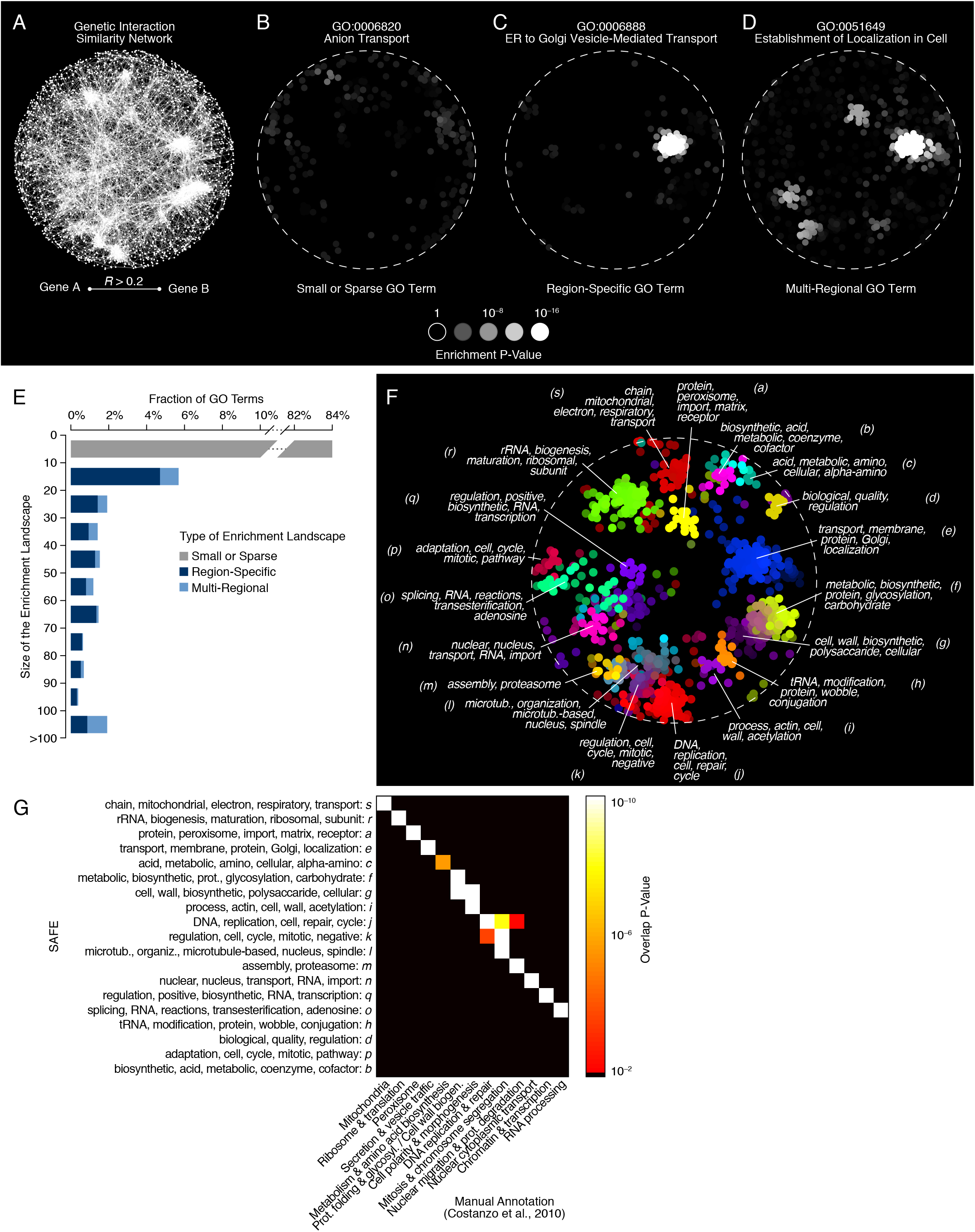
Annotating the yeast genetic interaction similarity network with Gene Ontology (GO) biological process terms. For details, see Experimental Procedures. (**A**) A map of the network, containing 2,838 nodes and 10,016 edges, was originally constructed in Costanzo et al. (Costanzo et al., 2010). Two genes (nodes) were connected if their genetic interaction profiles showed a Pearson correlation coefficient (*R*) greater than 0.2. Nodes were organized in two-dimensional space by applying the spring-embedded layout algorithm in Cytoscape (Shannon et al., 2003; Cytoscape.org, 2016). (**B-D**) SAFE annotation of the network with 4,373 GO biological process terms showed that different GO terms had different enrichment landscapes. (**B**) Most GO terms (84%), including anion transport (GO:0006820), were poorly enriched in the network. (**C**) 12% of GO terms, including ER to Golgi vesicle-mediated transport (GO:0006888), presented a region-specific enrichment landscape. (**D**) 4% of GO terms, including establishment of localization in cell (GO:0051649), showed multi-regional enrichment landscapes. (**E**) Distribution of 4,373 GO biological process terms based on the size and the type of their enrichment landscapes. (**F**) SAFE constructed a functional map of the network by combining all region-specific GO terms into 19 functional domains (*a*-*s*) based on the similarity of their enrichment landscapes. Different colors represent different functional domains. Each domain is labeled with a tag list, composed of the five words that occur most frequently within the names of the associated GO terms. (**G**) Analysis of the overlap between the SAFE-based and the manual (Costanzo et al., 2010) annotations of the network. Significance of overlap was measured by Fisher’s exact test. The resulting p-values were –log_10_ transformed and represented as a heatmap. Only p-values < 0.05 are shown.

SAFE mapped local enrichment for 4,373 GO biological process terms, each associated with at least one yeast gene, and revealed a great variation in the size, shape and relief of GO term enrichment landscapes (**Figure 2B-D**). Specifically, the vast majority of GO terms (84%) were only enriched within the neighborhoods of 10 or fewer genes, indicating that these terms were too small or too sparsely distributed throughout the network to be informative about its functional organization (**Figure 2B, E**). The remaining GO terms were enriched in more than 10 neighborhoods, but differed in the spatial distribution of their enrichments: 12% of GO terms were region-specific as they displayed a single peak of enrichment within a single region of the network (**Figure 2C, E**), whereas 4% of GO terms were multi-regional with two or more peaks of enrichment in different network regions (**Figure 2D, E**). The presence of multiple peaks suggested that each of the 174 multi-regional GO terms (4% of 4,373) comprised several functionally distinct subgroups of genes that localized separately within the network. Notably, each of these functional subgroups seemed to be well described by at least one of the 506 region-specific GO terms (12% of 4,373) which, all together, covered all multi-regional enrichment landscapes. Given that region-specific GO terms were sufficient to annotate the entire network, sparse and multi-regional terms were excluded from further analyses. Interestingly, while region-specific GO terms were generally smaller than multi-regional terms (p-value = 10^−49^, rank-sum test), their size distributions were largely overlapping (**Supplementary Figure 1**), suggesting that term size is not the sole responsible for landscape differences (see Discussion).

Since many of the 506 region-specific GO terms had overlapping enrichment landscapes and appeared to be functionally related, their contributions towards annotating the genetic interaction similarity network could be considered redundant. To minimize redundancy and simplify the annotation process, SAFE grouped the terms into 19 functional domains based on the similarity of their enrichment landscapes (Experimental Procedures). These domains, represented by different colors and labeled with their individual tag lists, formed a comprehensive GO-based functional annotation map of the genetic interaction similarity network (**Figure 2F**).

By comparing the automated functional map, produced by SAFE, to the manual annotation, carried out in (Costanzo et al., 2010), I found that the outcomes of the two approaches were highly consistent (**Figure 2F-G**). Specifically, all manually annotated regions significantly overlapped at least one of the 19 SAFE domains (all p-values < 2 × 10^−4^, Fisher’s exact test; **Figure 2G**). Furthermore, the GO terms associated with each domain were in agreement with the label assigned by manual analysis (**Figure 2G**). For example, SAFE recognized that domain *r*, previously associated with ribosome and translation-related processes, is enriched for 19 closely related GO terms, including “ribosome biogenesis” (GO:0042254) and “cytoplasmic translation” (GO:0002181). The complete list of GO terms associated with each functional domain is reported in **Supplementary Data 1**.

Interestingly, several network regions that were joint under a single functional label during manual analysis (Costanzo et al., 2010) were recognized as separate functional domains by SAFE (**Figure 2F-G**). For example, SAFE subdivided the “mitosis and chromosome segregation” region into two separate domains: one enriched primarily for mitosis and cell cycle-related GO terms (domain *k*) and the other for microtubule-based processes and cytoskeleton organization (domain *l*) (**Figure 2F-G**). In addition, SAFE refined the localization of metabolism-related genes (domains *b* and *c*) and recognized three previously unannotated network regions that were likely missed during manual analysis due to their reduced size and specific localization (domains *d*, *i* and *p*) (**Figure 2F-G**). These regions corresponded to the vacuolar H+-ATPase protein complex (domain *d*), the tRNA wobble base modification pathway (domain *i*) and the pheromone-induced cell cycle arrest factor (FAR complex, domain *p*).

### SAFE is robust against stochastic variations in network layout and differences in distance measures

To annotate the yeast genetic interaction similarity network, SAFE defined local neighborhoods using a distance metric that relies on the network map and thus may be sensitive to its non-deterministic nature. Specifically, two nodes belong to the same neighborhood if the shortest path between them, defined as the minimum distance one had to travel to get from one node to the other, was shorter than a chosen threshold (**Figure 1B**; Experimental Procedures). The lengths of the edges composing the paths were determined by the spring-embedded network layout implemented in Cytoscape (Cytoscape.org, 2016), which positions each node based on the balance of attractive and repulsive forces exerted, respectively, by nodes that are connected and disconnected from it (Experimental Procedures). Since the initial node positions are seeded randomly, the layout is non-deterministic and converges onto different final positions at every use (**Figure 3A-B, i**). This can potentially reshape local neighborhoods and change their enrichments, thus calling into question the accuracy of the resulting functional annotations.

**Figure 3.**
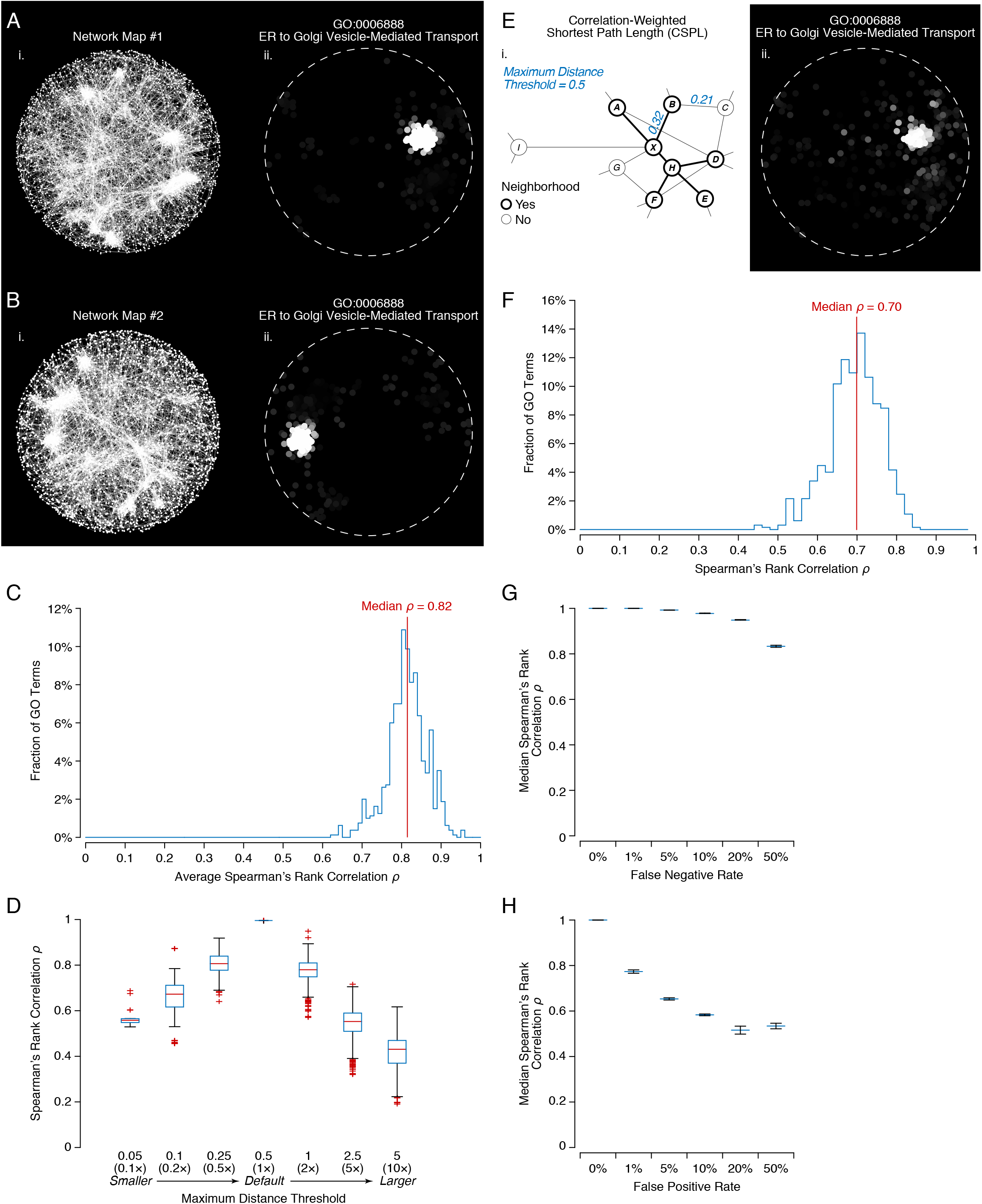
SAFE is robust against stochastic variations in network layout, differences in distance measures and noise in functional annotation standards. (**A-B**) An example of two different network maps produced by two independent runs of the spring-embedded network layout (i) and their corresponding enrichment landscapes for a region-specific GO term (ii). (**C**) Distribution of correlations between GO term enrichment landscapes across different network maps. Ten network maps were generated by repeatedly applying the spring-embedded layout algorithm in Cytoscape (Cytoscape.org, 2016). Each map was re-annotated with 4,373 GO biological process terms. For each GO term, Spearman’s rank correlation ρ was computed for all 45 pairs of enrichment landscapes (10 × 9 / 2 = 45), excluding the ones that had fewer than 10 enriched neighborhoods. The resulting ρ values were grouped by GO term, averaged and plotted as a distribution. (**D**) Distribution (median, 1^st^-3^rd^ quartiles ± 1.5× the range between the quartiles) of correlations between GO term enrichment landscapes across different neighborhood size thresholds. The maximum distance threshold (MDT), used to define neighborhood membership, was varied from the 0.05^th^ to the 5^th^ percentile of all pair-wise distances in the network. At each variation, the enrichment landscapes of all 4,373 GO terms were re-calculated and compared to the default landscape (MDT = 0.5^th^ percentile) using Spearman’s rank correlation, if both landscapes presented at least 10 enriched neighborhoods. (**E**) Definition of correlation-weighted shortest path length (CSPL) (i) and an example its corresponding enrichment landscape (ii). According to CSPL, the length of every path step equals to 1 minus the correlation of the genetic interaction profiles of the two connected genes. (**F**) Distribution of correlations between GO term enrichment landscapes computed using different distance metrics (MSPL vs CSPL). The network map (A) was re-annotated using the CSPL distance metric and the resulting enrichment landscapes were compared to their MSPL-derived versions using Spearman’s rank correlation, provided that both landscapes had at least 10 enriched neighborhoods. (**G-H**) Correlation of GO term enrichment landscapes before and after the introduction of false negative (G) or false positive (H) annotations. Each GO term was systematically altered by randomly introducing 1%–50% of false negative or false positive annotations. For false negatives, the indicated fraction of true positive annotations was set to negatives. For false positives, the indicated fraction of true negative annotations was set to positives. Each alteration was performed 1,000 times. After each alteration, GO term enrichment landscapes were recomputed and compared to their error-free versions using Spearman’s rank correlation, provided that both landscapes had at least 10 enriched neighborhoods. The median of all GO term correlations was calculated after each alteration. The medians were then averaged across 1,000 trials and plotted as a function of false negative (G) and false positive rates (H). Error bars represent standard deviations of the 1,000 medians.

To assess the robustness of SAFE to stochastic variations in network layout, I generated 10 different maps of the genetic interaction similarity network by repeatedly applying the spring-embedded layout (**Figure 3A-B, i**). I used SAFE to annotate each map with GO biological process terms and compared the enrichment landscapes of each individual term across all 45 map pairs (10 × 9 / 2 = 45) (**Figure 3A-B, ii**). I found that GO terms enriched within the neighborhoods of at least 10 genes showed highly similar enrichment landscapes in any two network maps (median Spearman’s rank correlation *ρ* = 0.82; **Figure 3C**). This indicates that, despite differences in absolute node positioning, the neighborhoods were largely unchanged and tended to be enriched for the same GO terms to the same degree. Importantly, the size of the enrichment landscapes was also consistent: of all GO terms that had at least 10 enriched neighborhoods in one map, 83% made the same threshold in at least 5 maps and 67% in all 10 (**Supplementary Figure 2A**).

Besides node positioning, the enrichment of local neighborhoods may also be affected by the maximum distance threshold that defines neighborhood membership (**Figure 1B**). By default, this threshold is set to the 0.5^th^ percentile of all pair-wise distances in the network (Experimental Procedures); however, higher or lower thresholds can also be chosen to produce larger or smaller neighborhoods. To determine whether neighborhood size affects the enrichments, I varied the distance threshold over a range of values from 0.1 to 10 times the default setting and, at each step, re-computed the enrichment landscapes for all GO biological process terms. I found that distance thresholds within a factor of two (0.5 and 2 times the default value) produced highly similar enrichment landscapes (median Spearman’s rank correlation *ρ* = 0.81 and *ρ* = 0.78, respectively; **Figure 3D**) and caused limited variation in the set of GO terms with more than 10 enriched neighborhoods (**Supplementary Figure 2B**). This indicates that neighborhoods are fairly robust to changes in size and, within a two-fold range, the choice of a distance threshold is not critical for mapping enrichment (Discussion).

Since neither the absolute node positioning nor the specific distance threshold appeared to have a significant impact on GO term enrichment landscapes, I asked whether the definition of distance itself also be considered a secondary parameter. To test this hypothesis, I chose an alternative distance metric and measured how much it affects neighborhood enrichments. In the genetic interaction similarity network, distance between two genes can be measured using the correlation-weighted shortest path length (CSPL). According to this metric, the length of each step equals 1–*R*, where *R* is the correlation of the genetic interaction profiles of the two connected genes (**Figure 3E, i**). Unlike the default map-weighted shortest path length (MSPL), the correlation-based metric is map-independent because *R* values were not used by the layout algorithm to generate the network map: the network was binarized at the *R* = 0.2 threshold and all edges above the threshold were assigned equal weight (Experimental Procedures). Using CSPL, I re-defined local neighborhoods and re-calculated their enrichments for GO biological process terms. For each individual GO term, I compared the enrichment landscapes produced by the MSPL and the CSPL distance metrics (**Figure 3A, ii**; **Figure 3E, ii**) and found them to be very similar (median Spearman’s rank correlation *ρ* = 0.70; **Figure 3F**). This indicates that distance definition is also not critical for mapping local enrichments and the neighborhoods defined with or without a network map tend to be functionally equivalent.

### SAFE is robust against moderate noise in the functional annotation standard

In addition to variations in network layout and distance metrics, the accuracy of network annotation may also depend on the quality of the functional annotation standard. Even functional resources as authoritative as GO are not free from annotation errors and may, in theory, produce spurious or incomplete local enrichments that could distort the overall view of the network’s functional organization. To test the robustness of SAFE to annotation noise, I systematically altered all GO biological process terms by randomly introducing false positive or false negative annotations at varying rates (from 1 to 50% of all annotations). After each alteration, I used the modified terms to re-annotate the genetic interaction similarity network and compared the resulting enrichment landscapes to their error-free versions. From this comparison, I found that SAFE is extremely robust to missing annotations (**Figure 3G**): a false negative rate (FNR) as high as 20% failed to significantly affect the enrichment landscapes of most GO terms (median similarity before and after alteration *ρ* = 0.95; Spearman’s rank correlation). In contrast, false positive annotations had a much larger impact (**Figure 3H**): the enrichment landscape of a GO term with a false positive rate (FPR) of 1% showed a median similarity of *ρ* = 0.78 (Spearman’s rank correlation) to its error-free version. Importantly, while both false positive and false negative annotations generally decreased the number of GO terms enriched within the neighborhoods of at least 10 genes (by 20% and 27% at FNR = 20% and FPR = 1%, respectively), the introduction of neither type of error caused an appearance of new, previously unobserved, GO term enrichments (**Supplementary Figure 2C-D**).

### SAFE annotates networks with chemical genomic data and recapitulates known drug modes-of-action

The automated GO-based annotation of the genetic interaction similarity network proved to be accurate (**Figure 2**) and robust to numerous sources of variation (**Figure 3**). However, annotating a network with a single type of biological information, such as GO, is unlikely to provide a full picture of the network’s functional organization because of the inherent biases and limitations of all functional standards. A more valid strategy is to use multiple independent sources of functional data and apply them iteratively to annotate the same network. Such an approach would not only provide a more realistic description of the network, but could also reveal unexpected relationships between functional standards.

In yeast, a rich source of genome-wide functional data is provided by chemical genomics, a powerful technology for characterizing gene functions and identifying the molecular modes-of-action of chemical compounds (Ho et al., 2011; Roemer et al., 2012). In a chemical genomic screen, a genome-wide collection of yeast mutants is grown in the presence of a chemical compound and the relative fitness of each mutant is measured with respect to an untreated control (Giaever et al., 2002). Identifying mutants that are particularly resistant or sensitive to a given compound is instrumental for mapping out pathways that mediate the compound’s toxicity or are required to protect the cell against its detrimental effects (Parsons et al., 2004; Parsons et al., 2006; Hillenmeyer et al., 2008; Hoepfner et al., 2014; Lee et al., 2014). I hypothesized that SAFE could assist the identification of these pathways by annotating the genetic interaction similarity network with chemical genomic data. Specifically, I predicted that SAFE could identify network regions that are enriched for genes that confer sensitivity or resistance to a given compound and determine whether the same regions are enriched for GO biological process terms that would be indicative of the compound’s mode-of-action.

To test this prediction, I used one of the most recent chemical genomics datasets in *S. cerevisiae*, which measured quantitative fitness scores for ∼5,000 homozygous deletion mutants upon exposure to 132 chemical compounds with known modes-of-action (Hoepfner et al., 2014). For each compound, low or high fitness scores indicated mutants that grew slower or faster compared to their corresponding untreated controls and the median of the population (Hoepfner et al., 2014). Using the genetic interaction similarity network and 132 sets of fitness scores, SAFE generated 132 compound-specific fitness enrichment landscapes that mapped the relative distribution of sensitive and resistant mutants throughout the network (**Figure 4A-B**).

**Figure 4.**
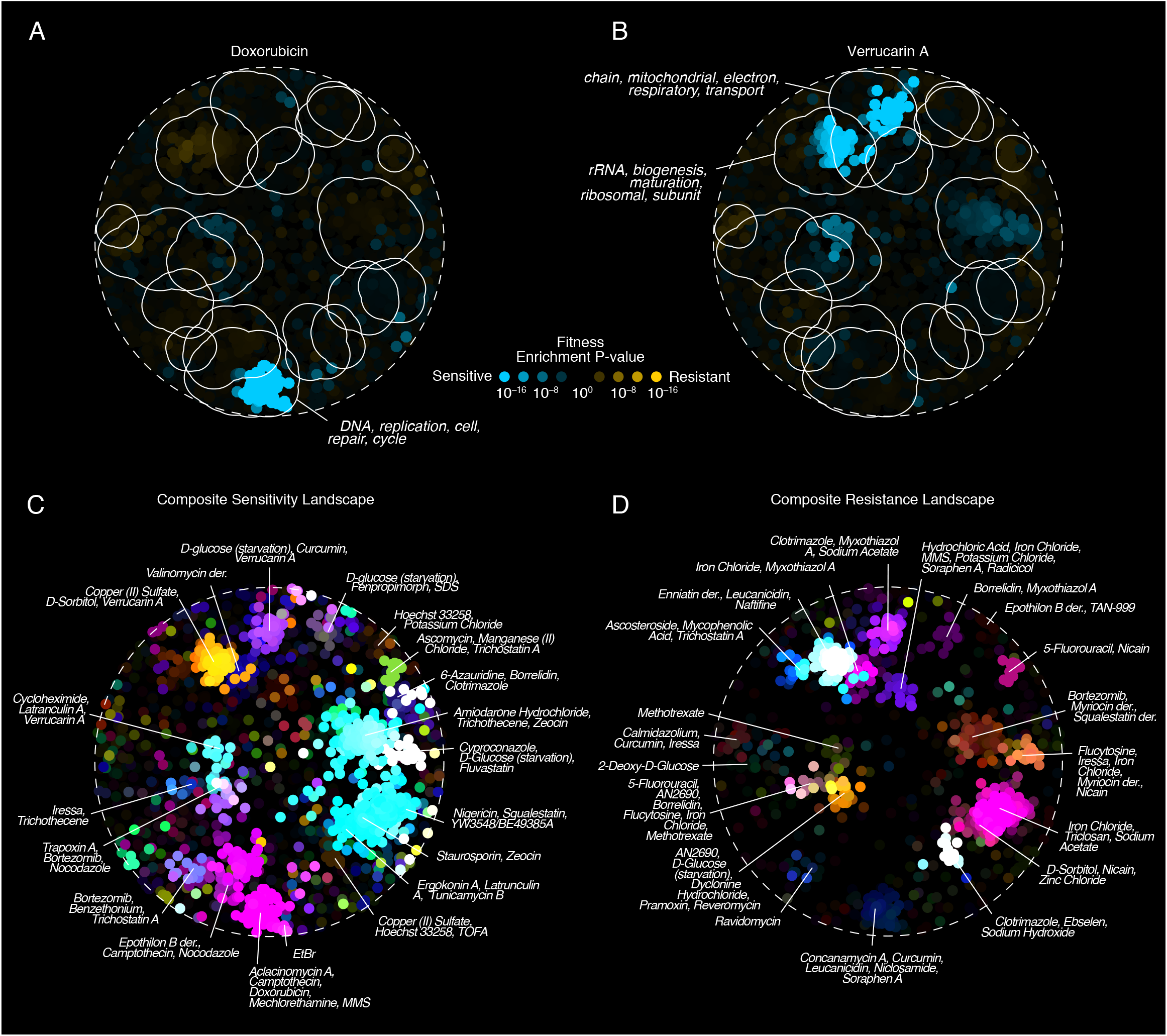
SAFE annotates the genetic interaction similarity network with chemical genomic data and recapitulates the modes-of-action of known drugs. (**A-B**) A fitness enrichment landscape maps local enrichment for genes associated with sensitivity or resistance to a chemical compound. For reference, the outlines of the 19 GO-based functional domains (Figure 2F) are shown. (**A**) Fitness enrichment landscape of doxorubicin, a DNA intercalator that blocks the progression of topoisomerase II and prevents DNA replication. (**B)** Fitness enrichment landscape of verrucarin A, a protein synthesis inhibitor with a known toxicity towards mitochondria. (**C-D**) The global map of chemical genomic annotations of the genetic interaction similarity network. Chemical compounds with similar sensitivity (C) or resistance (D) enrichment landscapes were assigned similar colors and summed proportionally to their relative enrichment within each neighborhood. Three examples and the total number of compounds enriched in each region are listed. Both region-specific and multi-regional chemical compounds were included. For reference, the outlines of the 19 GO-based functional domains (Figure 2F) are shown.

By examining the compound-specific landscapes (**Figure 4A-B**) in the context of the GO biological process map (**Figure 2F**), I found them to be highly consistent with our current knowledge about the compounds’ modes-of-action. For example, mutants sensitive to doxorubicin, a DNA intercalator that prevents DNA replication by blocking the progression of topoisomerase II (Momparler et al., 1976; Fornari et al., 1994; Tacar et al., 2013), were specifically enriched within the network region associated with DNA replication and repair GO terms (**Figure 4A**). Similarly, regions enriched for ribosome-and mitochondria-related GO terms were also overrepresented for mutants sensitive to verrucarin A, a protein synthesis inhibitor (Hernandez and Cannon, 1982) with a reported toxicity towards mitochondria (Schappert and Khachatourians, 1986) (**Figure 4B**).

Using the full set of 132 fitness enrichment landscapes, SAFE generated a global map of chemical genomic annotations for the genetic interaction similarity network (**Figure 4C-D**). This map grouped chemical compounds based on the similarity of their cellular effects (sensitivity and resistance analyzed separately) and recapitulated general trends that have long been associated with the yeast response to chemical treatment (**Figure 4C-D**).

For example, several studies reported that mutations affecting intracellular transport sensitize the cell to a wide range of chemical compounds and make up the most significant functional category among the multi-drug resistance genes in the yeast genome (Parsons et al., 2004; Hillenmeyer et al., 2008; Zakrzewska et al., 2010). Vesicle-mediated transport controls the localization of membrane proteins, including drug transporters and signaling receptors, and thus is expected to play a central role in regulating the efflux/influx of substances from/into the cell. Consistent with these observations, SAFE showed that the network region enriched for intracellular localization, transport and secretion GO terms was also the most common site for chemical sensitivity enrichment: 62 of the 132 compounds (47%) showed sensitivity landscapes that overlapped with the region (**Figure 4C**, **Supplementary Data 2**).

Similarly, resistance to multiple chemical treatments has been previously linked to the inactivation of ribosomal gene function (Zakrzewska et al., 2010). The growth benefit conferred by inactive ribosomes during chemical stress is consistent with the downregulation of ribosomal gene expression as part of the environmental stress response (ESR) program (Warner, 1999; Holcik and Sonenberg, 2005). The ESR program enables the cell to cope with different types of exogenous perturbations by activating a general defense strategy that includes the deceleration of protein synthesis and the delay of cell cycle progression (Gasch et al., 2000; Causton et al., 2001; Gasch and Werner-Washburne, 2002). Consistent with the nonspecific nature of this response, SAFE showed that the most common resistance enrichment landscape, characteristic of 45 of the 132 compounds (34%), involved the network region enriched for ribosome biogenesis and translation functions (**Figure 4D**, **Supplementary Data 2**).

### SAFE provides novel insights into the molecular basis of resistance to bortezomib

By systematically examining a biological network and mapping local enrichment for sensitivity and resistance to chemical compounds, as well as for GO term annotations, SAFE provides a comprehensive view of the cellular response to chemical treatment. In doing so, SAFE not only displays known modes-of-action in a simple and intuitive visual form, but can also uncover new response patterns and improve our understanding of chemical bioactivity. To illustrate this potential application of SAFE, I examined the fitness enrichment landscape of bortezomib, a proteasome inhibitor approved for treating multiple myeloma and mantle cell lymphoma and undergoing clinical trials for several other types of cancer (Chen et al., 2011) (**Figure 5**).

**Figure 5.**
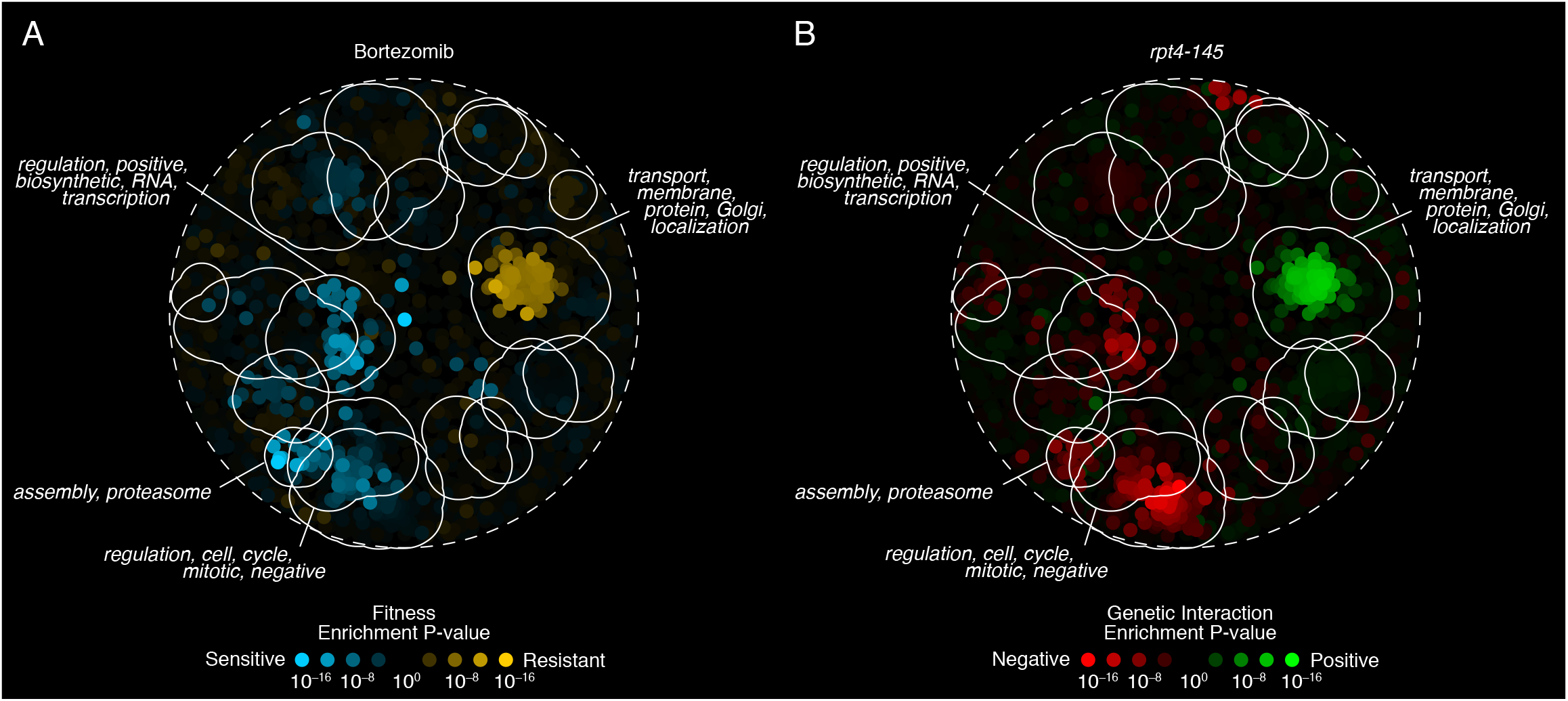
SAFE provides novel insights into the molecular basis of resistance to bortezomib. For reference, the outlines of the 19 GO-based functional domains (Figure 2F) are shown. (**A**) The fitness enrichment landscape of the proteasome inhibitor bortezomib shows that sensitivity to the drug is associated with mutations in proteasome-mediated protein degradation, cell cycle control and transcriptional regulation. In contrast, resistance to bortezomib is specific to genes involved in secretion and vesicle-mediated transport. (**B**) The genetic interaction enrichment landscape of *rpt4-145*, an essential member of the regulatory particle of the proteasome, is highly consistent with the fitness landscape of bortezomib and supports the role of secretory mutants in compensating for the effects of proteasome inactivation.

By blocking the proteasome’s proteolytic activity, bortezomib is thought to prevent the degradation of pro-apoptotic factors and indirectly promote programmed cell death in tumor cells (Chen et al., 2011). In addition, bortezomib’s anti-cancer effect was shown to increase upon addition of histone deacetylase inhibitors (Yu et al., 2003; Feng et al., 2007; Deleu et al., 2009; Canestraro et al., 2010), suggesting that transcriptional regulation may also play a role in mediating the drug’s activity. Consistent with these observations in human cells, SAFE showed that similar processes are likely impacted by bortezomib treatment in yeast: sensitivity to the drug was specifically overrepresented within regions of the genetic interaction similarity network that were associated with proteasome-mediated protein degradation, cell cycle control and transcriptional regulation (**Figure 5A**).

Intriguingly, SAFE also showed that resistance to bortezomib was strongly linked to the network region enriched for secretion and vesicle-mediated transport GO terms (**Figure 5A**). Although several reports have suggested that proteasome inactivation may cause endoplasmic reticulum (ER) stress, due to an accumulation of misfolded proteins in the ER (Lee et al., 2003; Obeng et al., 2006), it has not been anticipated that loss-of-function mutations in ER- or other vesicle-related functions could alleviate this or other proteasome-related stresses. A direct examination of bortezomib’s fitness data confirmed SAFE’s enrichment analysis: the four mutants with the highest resistance to the drug carried complete or partial deletions in genes regulating the formation, motility and fusion of vesicles traveling to and from the Golgi compartment – *YTP6*, *RIC1* and *RGP1* (Siniossoglou et al., 2000). In addition, several other proteins involved in Golgi-related transport were among the top 15 most resistant mutants.

Importantly, the statistical significance of this finding was also supported by a network-independent Gene Set Enrichment Analysis (GSEA) of bortezomib’s fitness data. GSEA determines whether members of a functional group tend to occur at the top or at the bottom of a ranked gene list and measures the probability of such distribution to arise by random chance (Subramanian et al., 2005). By applying GSEA to the ranked list of bortezomib’s fitness scores and all 4,373 GO biological process terms, I confirmed that processes such as “intra-Golgi vesicle-mediated transport” (GO:0006891) and “cytoplasm to vacuole targeting (CVT) pathway” (GO:0032258) are significantly associated with the most highly resistant mutants (p < 0.001 and p = 0.001, respectively; FDR < 0.05; **Supplementary Data 3**). Compared to SAFE, however, the primacy of these pathways in resisting bortezomib was much less apparent: the majority (58%) of the 48 significant GO terms detected by GSEA involved ion homeostasis, regulation of intracellular pH and other distantly related functions, associated with only moderately resistant mutants (**Supplementary Data 3**). Such a discrepancy indicates that SAFE’s fitness enrichment landscapes can uncover signals that are not easily detectable by other methods and can therefore provide unique insights into the action of chemical compounds.

To further validate the connection between vesicle-mediate transport and resistance to bortezomib, I asked whether, similarly to other drugs (Parsons et al., 2004; Costanzo et al., 2010), the effect of bortezomib could be mimicked by the genetic inactivation of its molecular target, i.e. the proteasome. Specifically, I hypothesized that mutations providing a relative growth advantage in the presence of bortezomib should also display a relatively high fitness when combined with proteasome mutants and should therefore result in positive genetic interactions.

To test this hypothesis, I obtained quantitative negative and positive genetic interactions for members of the core and the regulatory particles of the yeast proteasome (Experimental Procedures, **Supplementary Data 4**) and used them to annotate the genetic interaction similarity network with SAFE. For every gene encoding a proteasomal subunit, SAFE calculated a genetic interaction enrichment landscape which mapped local enrichment for the gene’s negative and positive interactors throughout the network (**Figure 5B**). I found that at least 7 of the 13 tested subunits of the proteasome regulatory particle were significantly enriched for positive genetic interactions within the vesicle transport region (**Supplementary Figure 3**). In particular, *rpt4-145*, a temperature-sensitive mutant in an essential ATPase of the regulatory particle that preferentially contributes to ER-associated protein degradation (ERAD) (Lipson et al., 2008), showed a genetic interaction enrichment landscape remarkably similar to bortezomib’s fitness enrichment landscape in both its negative and positive interaction enrichments (**Figure 5B**). It is important to note that a direct comparison between the fitness profiles of *rpt4-145* and bortezomib would also have revealed their similarity (*R* = 0.22, *p* < 10^-31^); however, determining what drives this similarity, i.e. specifically which positive and negative interactions are common to both profiles, could not have been accomplished by correlation analysis alone.

These findings support the hypothesis that mutations in secretory functions may partially compensate for proteasome inactivity and alleviate the damaging effects of bortezomib treatment, although the precise mechanisms of this alleviation are still unknown. Considering that resistance to bortezomib is a common complication in treating multiple myeloma and other cancers (Murray et al., 2014), understanding its molecular mechanisms is critical for designing targeted therapeutic approaches and effective drug combinations. The identification of secretory pathways as a potential focus of these new therapies illustrates the power of SAFE to uncover novel biological responses by comprehensively annotating biological networks with multiple independent functional standards.

### SAFE reveals functional structure in super-dense biological networks

By iteratively incorporating GO biological process, chemical genomic and genetic interaction annotations, SAFE produced accurate and informative functional maps of the genetic interaction similarity network. Despite its success, the annotation of genetic interactions may be considered a relatively easy task: the network shows a well-organized topology in which large modules of densely connected—and thus likely functionally related—nodes are easily distinguishable from one another, even by the naked eye (**Figure 2A**). In contrast, most other biological networks in yeast and other organisms have a more complex structure with higher edge density and lower node modularity (Jeong et al., 2001; Giot et al., 2003; Rual et al., 2005; Yu et al., 2008; Nayak et al., 2009; Arabidopsis Interactome Mapping, 2011). As a result, resolving the functional organization of these networks may be more difficult than suggested so far.

To assess whether SAFE can potentially annotate more complex biological networks, I first tested whether it could annotate a genetic interaction similarity network with a higher edge density. Given that, in the original network, two genes were connected if their profile correlation exceeded an arbitrary threshold (*R* > 0.2), I progressively reduced the stringency of the threshold (*R* > 0.18, *R* > 0.16 and *R* > 0.14) and increased the number of edges by 1.4, 2.1 and 3.4 fold (**Figure 6A, i**). As expected, the maps of the resulting three networks showed rapidly decreasing levels of visible structure and were nearly impossible to explore by eye (**Figure 6A, i**).

**Figure 6.**
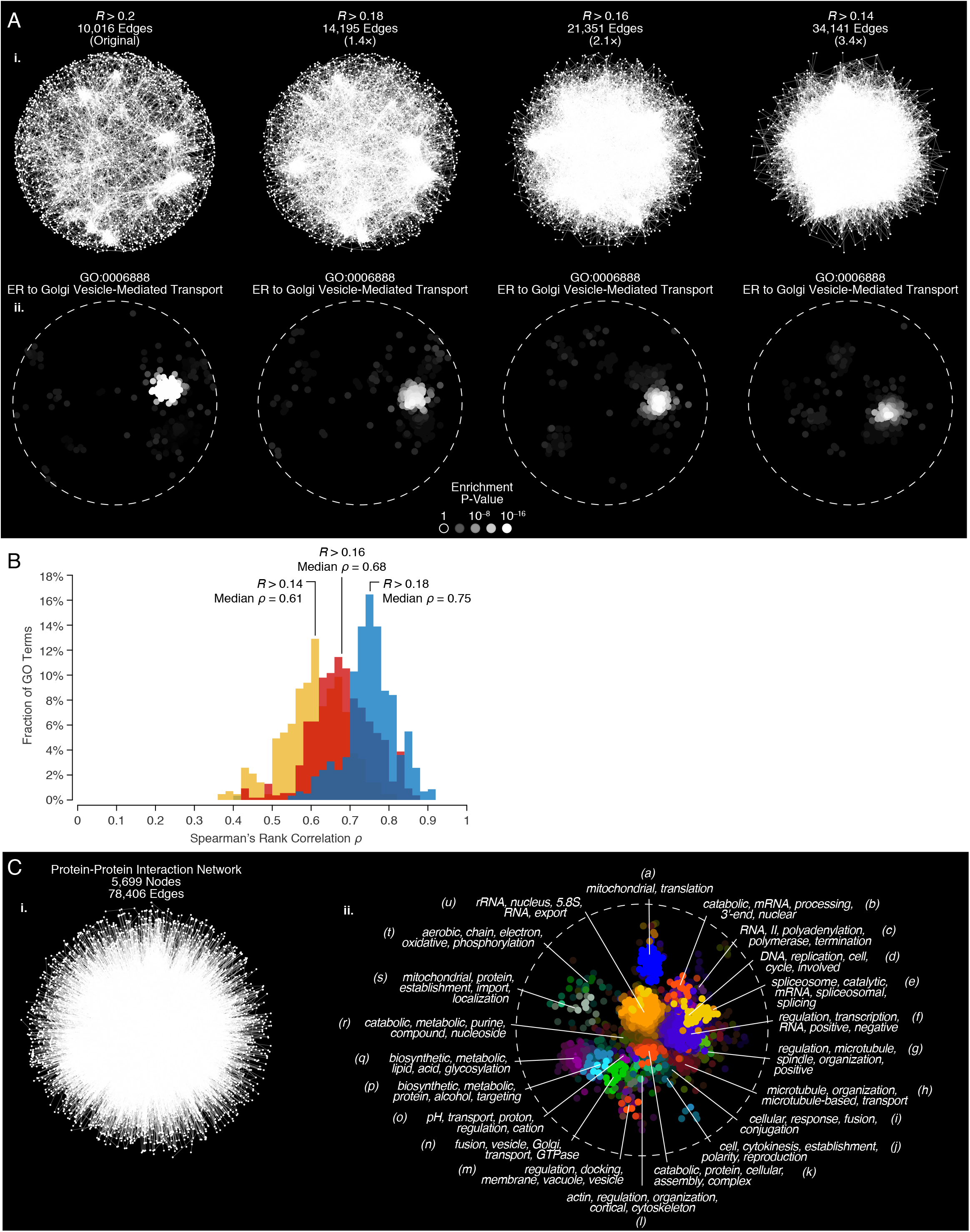
SAFE annotates dense genetic and protein-protein interaction networks. (**A**) **i**. Three additional versions of the genetic interaction similarity network were generated by progressively decreasing the correlation threshold (*R* > 0.18, *R* > 0.16 and *R* > 0.14). Compared to the original network (*R* > 0.2), the new networks had 1.4, 2.1 and 3.4 times more edges, respectively. The network maps were produced by applying the spring-embedded layout algorithm in Cytoscape (Cytoscape.org, 2016). **ii**. Despite the disappearing visible structure, SAFE detected functional enrichment in all three network maps. (**B**) Distribution of correlations between GO term enrichment landscapes across network densities. For every GO term, a Spearman’s rank correlation coefficient was computed between the original enrichment landscape, derived from the *R* > 0.2 network, and the enrichment landscapes derived from denser networks. Sparse and multi-regional GO terms were excluded from the analysis. (**C**) **i**. A global protein-protein interaction network, containing 5,699 nodes and 78,406 edges, was constructed from all yeast protein pairs sharing a physical interaction type in the BioGRID database (Chatr-Aryamontri et al., 2015) (Experimental Procedures). A map of the network was generated by applying the spring-embedded layout algorithm in Cytoscape (Cytoscape.org, 2016). **ii**. In a manner analogous to the genetic interaction similarity map (Figure 2), the protein-protein interaction map was annotated with SAFE using 4,373 GO biological process terms. All region-specific GO terms were combined into 21 functional domains (*a*-*u*) based on the similarity of their enrichment landscapes. Different colors represent different functional domains. Each domain is labeled with a tag list, composed of the five words that occur most frequently within the names of the associated GO terms.

I used SAFE to annotate the three network maps with GO biological process terms and found that local enrichment was easily detected in spite of the increasing edge density (**Figure 6A, ii**). Furthermore, the enrichment landscapes for each individual GO term were similar, albeit to a gradually decreasing extent, to the original landscape derived from the lowest density map (median Spearman’s rank correlations *ρ* = 0.75, *ρ* = 0.68 and *ρ* = 0.61; **Figure 6B**). Thus, even a ∼3-fold difference in the number of gene connections appeared to have a limited impact on the GO term enrichments in most gene neighborhoods.

The ability to map functional enrichment in a dense genetic interaction similarity network indicated that SAFE is sensitive to coherent functional structure. However, the existence of a functional structure itself may be a unique property of genetic interactions and may not be associated with any other biological network. To test whether that is the case, I examined the extensive yeast protein-protein interaction network which maps 78,406 physical binding events between 5,699 proteins in the cell (Chatr-Aryamontri et al., 2015) (Experimental Procedures). While protein-protein interactions are known to be highly nonrandom and to preferentially connect members of the same protein complex and other functionally related proteins (Cusick et al., 2005), a genome-wide map of the network, produced by the spring-embedded layout algorithm, showed no visible topological structure on a global scale (**Figure 6C, i**). To search for a potential functional structure, I used SAFE to annotate the network with GO biological process terms. Strikingly, SAFE revealed 21 large functional domains, each associated with a distinctive GO term enrichment landscape and a unique list of enriched GO terms (**Figure 6C, ii; Supplementary Data 5**). The resulting functional map strongly suggests that physical binding, just like genetic interaction profile similarity, assembles proteins into high level functional communities whose relative positioning within the network may provide new insight into the global functional organization of the yeast cell.

## DISCUSSION

Here I described the development, validation and application of SAFE, the first automated method for annotating biological networks and exploring their functional organization. Given a network and a visual map of its connectivity, SAFE localizes all network regions enriched for one or more functional attributes, such as Gene Ontology terms or quantitative phenotypes. In doing so, SAFE addresses three fundamental questions that help elucidating the network’s functional organization. First, are any regions of this network specifically associated with a given function or phenotype? Second, where in the network are these regions localized? And third, how does their localization compare to that of other functions or phenotypes? By answering these questions, SAFE builds a functional map of the network and enables the investigation of inter-process relationships within the cell.

Rapid and systematic identification of functional regions is a distinctive feature of SAFE that encourages the use of multiple complementary functional standards to annotate networks in an iterative manner. In this study, I showed that a sequential annotation of the yeast genetic interaction similarity network with GO biological process, chemical genomics and genetic interaction data provides a comprehensive view of the cellular response to chemical perturbation, linking genes and their functions to bioactive compounds and enabling the prediction of novel mechanisms of drug resistance (**Figure 4–5**).

Another distinctive feature of SAFE is that, through the power of visualization, it can improve our understanding of functional standards. For example, by annotating the genetic interaction similarity network with GO biological process terms, SAFE showed that some GO terms are enriched within a single network region, whereas others are multi-regional (**Figure 2B-E**). Although multi-regional GO terms tended to be generally larger than region-specific ones, term size alone could not fully explain the difference between the two groups (**Supplementary Figure 1**). An interesting possibility is that region-specific GO terms share a similar level of functional specificity, which is defined by the topology of the genetic interaction similarity network. If that is the case, these terms could be used to delineate a cross section of the GO hierarchy and form a flat subset of GO annotations, analogous to GO slim (Ashburner et al., 2000; Geneontology.org, 2016). Flat annotation resources have been instrumental for many genomic analyses thanks to their smaller size and lower redundancy (Myers et al., 2006; Costanzo et al., 2010). SAFE may offer the possibility to generate data-driven network-specific flat annotation standards that would enable more targeted functional analyses and simplify their biological interpretation.

A third feature that enables SAFE to annotate biological networks in an accurate and comprehensive manner is its relative robustness to incomplete or noisy data. Validation results showed that SAFE is largely insensitive to several types of input variation, including stochastic effects in network layout, different distance measures and overall density of the network (**Figures 3, 6**). In addition, moderate rates of false negative and false positive errors in functional annotation standards proved to have little impact on the outcome of SAFE (**Figure 3G-H**). Wrong or missing annotations are expected to scatter randomly throughout the network and fail to significantly alter the enrichments of local neighborhoods. This aspect of SAFE is particularly valuable for functional standards based on high-throughput or computationally inferred datasets where random noise, as well as systematic effects, are more likely to interfere with real functional relationships. For example, it has been reported that chemical genomics data may be affected by an experimental bias whereby deletion mutants constructed in the same laboratory share common secondary mutations and show similar growth phenotypes in response to a wide range of chemical compounds (Hoepfner et al., 2014). SAFE would not recognize these phenotypes as a reliable biological signal because genes mutated in the same laboratory do not preferentially interact with one another or localize to the same region of the genetic interaction similarity network (**Supplementary Figure 4**).

In addition to being robust to noise, SAFE is also sensitive to true biological signal, as shown by its ability to detect enrichment in super-dense genetic interaction similarity and protein-protein interaction networks (**Figure 6**). This feature enables SAFE to be applied to a wide range of biological networks and makes the method particularly beneficial to those that lack a visible topological structure on a global scale. Because of their complexity, these networks are often described as “hairballs” and considered inaccessible to biologists. However, SAFE may reveal that, behind an apparent disorderliness, there lies a functional structure that can be examined on a global scale without decreasing edge density and thus potentially loosing valuable biological information.

Finally, SAFE provides a unique opportunity to exhaustively examine the functional landscape of a network because it explores all local neighborhoods and does not focus exclusively on densely connected clusters. A neighborhood is a set of nodes located within a short distance from a source node, irrespective of the total number of connections between them. As a result, SAFE does not miss functional coherence within sparse network regions (**Figure 2F**) and is capable of annotating networks with virtually no variation in interaction density, such as a lattice. In this respect, SAFE is conceptually different from other approaches for network analysis, including clustering, and provides a more comprehensive overview of a network’s functional content.

Mapping the functional organization of biological networks is a complex task that requires accurate statistical analysis and informative data representations. While SAFE addresses some of the main challenges associated with this task, the method is still far from complete and presents vast opportunities for future development.

For example, the interpretation of the functional maps produced by SAFE would greatly benefit from a better understanding of network layout algorithms. Data-driven network layouts, such as the spring-embedded algorithm in Cytoscape (Cytoscape.org, 2016), are unsupervised methods for organizing nodes based on their connectivity. In its default settings, SAFE relies on layouts to identify local neighborhoods and visually map their functional enrichment. However, very little is known about how a particular layout should be chosen. Despite their great potential for uncovering hidden patterns within the data, layouts are typically used to generate esthetically pleasant network visualizations (Agapito et al., 2013) and are rarely the basis of any systematic network analysis. As a result, we have limited experience in evaluating network layouts and a poor understanding of their relative performance in the context of different networks.

So far, the decision to use the spring-embedded layout to represent and annotate the genetic interaction similarity network was founded on theoretical considerations: the basic working principles of the algorithm are in agreement with our understanding of genetic interactions and produce an easily interpretable outcome (Costanzo et al., 2010). While that decision has proved to be very successful, SAFE offers a unique opportunity to quantitatively support it or to identify a potentially better approach for annotating the network. To achieve this, SAFE must be systematically applied to annotate and evaluate different layouts for the same network using a common set of functional attributes. The ultimate outcome of this analysis would be the identification of an optimal layout for each individual network type and the establishment of a common ground for comparing different networks to one another.

Quantitative comparison of biological networks is indeed a major goal of systems biology (Sharan and Ideker, 2006). Without a deep understanding of how genes, pathways and processes are connected across different kinds of molecular networks, we have little hope of developing successful strategies for integrating networks into a single comprehensive model of a living cell. By mapping enrichment for the same functional attributes in different networks (**Figure 2**, **Figure 6**), SAFE can make an important contribution to this goal. However, rigorous statistical approaches must be implemented to compare SAFE maps across networks and draw meaningful conclusions about their differences and similarities.

In summary, SAFE represents an important step towards advanced understanding of biological networks. Unlike most other methods for network analysis, SAFE provides a global perspective onto the functional organization of a network by mapping statistical associations between functional groups and network regions. SAFE shows that, despite previous lack of attention, network layouts coupled with robust enrichment analysis are a valid strategy for studying molecular networks and gaining insight into the biological systems they represent. As both the number and the extent of molecular networks grow exponentially over time, understanding their functional organization with methods similar to SAFE is becoming of primary importance.

## Acknowledgements

I am deeply grateful to Dmitriy Gorenshteyn for his invaluable help with writing this manuscript. Also, I would like to thank Amanda Amodeo, David Botstein, Michael Costanzo, Charles Boone and Brenda Andrews for reading the manuscript and making useful suggestions. The Boone and Andrews labs also generously provided the proteasome genetic interaction data. This work was supported in its entirety by the Lewis-Sigler fellowship at Princeton University.

## EXPERIMENTAL PROCEDURES

SAFE is openly accessible as MATLAB code at https://bitbucket.org/abarysh/safe under the MIT license. I strongly encourage users to subscribe as “watchers” to the Bitbucket repository in order to receive live updates about new code releases and bug fixes.

The basic flow of the algorithm, including key inputs and outputs, is described below. A more detailed description, including examples, is provided in the README file of the Bitbucket repository.

### The SAFE algorithm

#### Software requirements

MATLAB, Bioinformatics Toolbox, Statistics and Machine Learning Toolbox

#### Input data

1) A network *G* = (*V,E*) that consists of a set of nodes *V* and a set of undirected edges *E* ⊆*V* ×*V*. Edge weights *w_e_* |*e* ∈*E* can be binary (*w_e_* ∈{0,1}) or quantitative (*w_e_* ∈*R*).
2) A functional annotation standard in the form of a matrix *A*= *L*× *F*, where *L* are node labels and *F* are functional node attributes. The matrix *A* can contain binary or quantitative functional attributes. In case of binary attributes (e.g., Gene Ontology annotations), the entry *a_ij_* in matrix *A* equals 1 if the *i*-th node is annotated to the *j*-th functional attribute, and 0 otherwise. In case of quantitative attributes (e.g., a chemical genomics dataset), the entry *a_ij_* in matrix *A* equals to the quantitative measurement associated with the *i*-th node in the *j*-th experimental condition.

#### Algorithm

1) Load the network.
2) Optional. Create a new network map by applying a force-directed network layout algorithm (e.g., Kamada-Kawai (Kamada and Kawai, 1989) or Fruchterman-Reingold (Fruchterman and Reingold, 1991)) implemented as part of the MatlabBGL toolbox (Gleich, 2009).
3) Load the functional annotation standard.
4) For every node *v_i_* in network *G*, define a local neighborhood as the set of nodes *U_i_* that can be reached from *v_i_* by traveling no more than the maximum distance *d*. Distance between any two nodes can be measured as an unweighted, a map-based (default) or an edge weight-based shortest path length between them.

a. In the unweighted case, traversing the edge between nodes *v_i_* and *v_k_* has a constant cost *C_ik_* = 1.
b. In the map-based case, traversing the edge between nodes *v_i_* and *v_k_* has the cost of the Euclidean distance between the nodes on the network map: 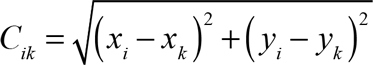 where *x* and *y* are the spatial coordinates of nodes *v_i_* and *v_k_*.
c. In the edge weight-based case, traversing the edge between nodes *v_i_* and *v_k_* has the cost of the original edge weight *w_ik_* (see Input data, above). By default, the maximum distance threshold *d* equals to the 0.5^th^ -percentile of all pair-wise node distances in the network.
5) Calculate the enrichment of the neighborhood *U_i_* for each functional attribute *F_j_ ∈ F* in the annotation standard *A*. Two ways of calculating enrichment are used, depending on whether the attributes are binary or quantitative.

a. For binary attributes, the enrichment is calculated by performing a one-tailed Fisher’s exact test. The significance of the enrichment is determined by the probability *p_ij_* that a random overlap between *U_i_* and *F_j_* will fall in the interval [*S_ij_* ∞):

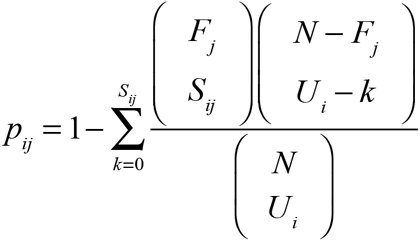

where: *N* = number of nodes in the network *F_j_* = number of nodes in the network annotated to functional attribute *F_j_* *U_i_* = number of nodes in neighborhood *U_i_* *S_ij_* = number of nodes in neighborhood *U_i_* annotated to functional attribute *F_j_*
b. For quantitative attributes, the enrichment is calculated using an empirical test. A neighborhood score *S_ij_* is computed by summing the quantitative attribute values *F_j_* for all nodes in *U_i_*. The score *S_ij_* is then compared to the mean µ and the standard deviation σ of 1,000 random neighborhood scores obtained by reshuffling the network labels and, consequently, the functional attribute values. The significance of the enrichment is determined as the probability that a single observation from a normal distribution with mean µ and standard deviation σ will fall in the interval (–∞ *S_ij_*] or [*S_ij_* ∞), depending on whether the highest or the lowest scores are of interest.
6) Convert neighborhood significance p-values *p_ij_* into neighborhood enrichment scores *O_ij_*, normalized to a range from 0 to 1, by computing:

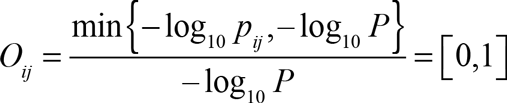

where *P* = 10^−16^ and corresponds to the smallest p-value that Matlab can calculate in a Fisher’s exact test. A neighborhood *U_i_* is considered significantly enriched for the functional attribute *F_j_* if:

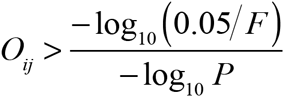

where *F* is the number of functional attributes in the annotation standard. This is equivalent to a Bonferroni multiple testing correction of the enrichment p-values *p_ij_*. The enrichment landscape of the functional attribute *F_j_* is defined as the vector of *O_j_* values for all nodes *i* in the network.
7) Optional. Plot the enrichment landscapes for all or some of the functional attributes individually. Examples are Figure 2B-D, Figure 4A-B and Figure 5A-5B.
8) Optional. Restrict the analysis to region-specific functional attributes using one of the three available methods.

a. Hartigan’s dip test of unimodality (Hartigan and Hartigan, 1985): defines a functional attribute as region-specific if the distribution of pair-wise distances between its significantly enriched neighborhoods is unimodal.
b. Radius-based method (default and fastest): defines a functional attribute as region-specific if at least 65% of significantly enriched neighborhoods are within a distance of 2*d*, where *d* is the radius of the neighborhood (see #4 above).
c. Subtractive clustering-based method (Chiu, 1994; Yager and Filev, 1994): defines a functional attribute as region-specific if there is at most one cluster center among the significantly enriched neighborhoods.
9) Optional. Group functional attributes into functional domains based on the similarity of their enrichment landscapes. Distance between functional attributes *F_j_* and *F_k_* is computed using one minus Jaccard (default) or Pearson correlation coefficients between their corresponding enrichment landscapes *O_j_* and *O_k_*. Given an *F* × *F* distance matrix, functional domains are formed using agglomerative hierarchical clustering with average linkage. Domains are defined by a distance threshold, set by default at 75% of the cluster tree height.
10) Assign a random RGB color to each functional domain. All functional attributes in the same domain are assigned the same color.
11) Determine the color *RGB_i_* of every node *v_i_* in the network *G* by computing a weighted average of the colors associated with the functional attributes for which the neighborhood *U_i_* is significantly enriched. The weights correspond to the square of the neighborhood enrichment scores *O_ij_* for each functional attribute *F_j_*.

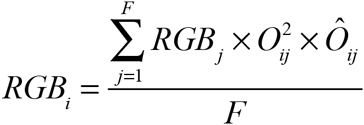

where *Ô_ij_* equals 1 if the neighborhood is significantly enriched for functional attribute *F_j_* and 0 otherwise.
12) Output the results.

a. Plot the composite functional map of the network. Examples are Figure 2F, Figure 4C-D and Figure 6C (ii).
b. Generate an automatic label for each functional domain by identifying the five most recurrent words in the denominations of the functional attributes that belong to that domain.
c. Print all data, including the list of functional domains, their automatically generated labels and the complete list of functional attributes enriched within each domain, into text files.
13) Save all data and settings into a SAFE session file.

### Annotation of the yeast genetic interaction similarity network

The yeast genetic interaction similarity network was constructed as described in Costanzo et al. (Costanzo et al., 2010). Briefly, similarity of genetic interaction profiles for all pairs of 1,712 query genes and all pairs of 3,885 array genes was computed using Pearson correlation coefficients (*R*). Correlation values for gene pairs that have been tested both as queries and as arrays were averaged. Gene pairs presenting similarities greater than *R* = 0.2 were connected in a network and visualized using the edge-weighted spring-embedded layout in Cytoscape 2.8 (Shannon et al., 2003), which corresponds to an implementation of the Kamada-Kawai layout algorithm (Cytoscape.org, 2016). Despite the choice of an edge-weighted version of the algorithm, it was later discovered that a bug in Cytoscape 2.8 caused edge weights to be ignored. Thus, running the edge-weighted layout algorithm was equivalent to binarizing the network at the *R* = 0.2 threshold and running the unweighted version of the algorithm. The bug was fixed in Cytoscape 3.0.

The Gene Ontology (GO) biological process data and the yeast gene association files were downloaded from www.geneontology.org on August 19, 2014 (Ashburner et al., 2000). Annotations were propagated from child to parent terms, such that a gene was associated with a GO term if it was directly annotated to the term or any of its descendants.

The chemical genomics dataset from Hoepfner et al. (Hoepfner et al., 2014) was downloaded from the Dryad digital repository (Hoepfner D, 2013) on December 3, 2013. Only homozygous profiling (HOP) data for 132 chemical compounds with known modes of action (as listed in the Supplementary Table 1 of that publication) were used.

The quantitative genetic interaction data involving 13 members of the proteasome regulatory subunit were obtained and processed as described previously (Baryshnikova et al., 2010a; Baryshnikova et al., 2010b). Specifically, the laboratories of Charles Boone and Brenda Andrews (University of Toronto) conducted genome-wide synthetic genetic array (SGA) experiments to construct double mutants involving a deletion or a temperature-sensitive allele of a proteasome member (*RPN1*, *RPN5*, *RPN6*, *RPN7*, *RPN10*, *RPN11*, *RPN12*, *RPT1*, *RPT2*, *RPT3*, *RPT4*, *RPT6* and *SEM1*) and the deletion mutants of all non-essential genes in the yeast genome. Quantitative double mutant fitness values were measured using colony size and compared to the fitness values of the two corresponding single mutants to produce a genetic interaction score that quantifies negative and positive deviations of the observed double mutant fitness from the expected combination of the two singles. The genetic interaction scores for these 13 SGA screens are available as **Supplementary Data 4**.

### Annotation of the yeast protein-protein interaction network

A complete set of yeast protein-protein interactions was downloaded from BioGRID on April 26, 2015. The dataset was filtered to include only “physical” interaction types and exclude “biochemical activity” and “protein-RNA” experiments. The network was visualized using the spring-embedded layout in Cytoscape 3.2 (Shannon et al., 2003; Cytoscape.org, 2016).

